# Lithographic structuring of thermoresponsive hydrogel on a micron scale

**DOI:** 10.1101/2025.01.29.635555

**Authors:** Katharina Esch, Katja Zieske

## Abstract

Hydrogel microstructures and microchannels are important tools in biophysical bottom-up approaches for studying the interactions between extracellular obstacles or boundaries and cells. In prior studies, several microstructuring techniques, cell sources, and gel types have been employed to mimic static cellular environments. However, the dynamic nature of the extracellular environment, which exerts forces on cells, plays an important role in tissue-level contexts. To date, only few approaches have been implemenmted to mimic these forces of the extracellular environment.

Here, we present an approach for generating arrays of hydrogel microstructures, that expand and contract dynamically in response to temperature modulations. To do this, we use a thermoresponsive hydrogel and a custom-built lithographic setup. Through direct lithographic photopatterning of a liquid phase, we fabricate smart hydrogel microstructures at the micrometer scale (10-100 μm) with well-defined geometries. These hydrogel microstructures display reversible size changes of over 70% triggered by temperature modulations.

Our study represents a step towards mimicking the dynamic properties of biological boundaries and obstacles. The dynamic nature of smart hydrogel microstructures holds the potential to mimic the micron-scale dynamics of extracellular environments. Our results may thus be relevant for the fields of bioengineering and systems level biophysics offering opportunities for manipulating and exploring cellular interactions with dynamic obstacles.

## Introduction

Dynamic rearrangements of biological matter play significant roles across various scales, influencing the organization of organisms, tissues, individual cells, and molecular structures. At the tissue-level, cells constantly navigate an environment characterized by dynamic factors, such as rhythmic muscle movements, blood flow, and intercellular interactions.To study these dynamic processes between cells and the extracellular environment, bioengineered environments created using bottom-up approaches offer promising tools. However, current approaches fail to fully replicate the whole complexity and dynamics of the extracellular environment.

Progress in culturing 3D multicellular models and embedding cells in soft polymer environments provided important insights into cell physiology and behavior under conditions that mimic natural environments more closely as compared to experiments on glass or polystyrene substrates (1,2), but it remains a challenge to probe the influence of dynamic obstacles and boundaries. Therefore, the question arises whether dynamic microstructures with varying geometries can be generated at the scale of 10-100μm to mimic the dynamics of cells and minimal tissue contexts.

Hydrogels are polymer networks with a high water content that mimic certain aspects of the extracellular environment (3). Some widely used polymers include polyethylene glycol (PEG), polyacrylamide, and biological polymers such as collagen (4) and DNA (5). While static hydrogels are frequently applied as an artificial extracellular environment (6), the need for dynamic, micro-structured hydrogels arise for replicating dynamic changes in cellular environments. Dynamic systems, such as cyclic strain bioreactors, which stretch the entire substrate, and consequently the attached cells, have been developed. (7–10). More recently, stimulus-responsive hydrogel platforms for stretching single adherent cells have been engineered (11–15). These approaches are promising for implementing spatio-temporally controlled dynamics in extracellular environments. Thereby, hydrogels that respond to changes in temperature by changing their size are promising materials for creating dynamic micron scale structures. Poly(N-isopropylacrylamide) (NiPAAm) is a thermoresponsive polymer with a critical temperature of 32°C (16). Above the critical temperature NiPAAm is coiled up, while below it, the polymer transitions into extended chains (17). This property has been employed in surface coating and hydrogel applications in biology and biomedicine. For instance, NiPAAM substrates have been used to induce detachment of adherent cells upon cooling below the critical temperature (18–21). NiPAAm hydrogels have also been employed in the context of release and delivery of drugs (22) and for engineering micro-robots (23). In addition, NiPAAm based substrates enabled the local stretching of cells and localized volume change of the substrate upon stimulation (11–14). Thus, for designing micron scale structures with integrated dynamics upon temperature changes and subsequently, for generating systems that expose cells to micromechanical forces in their environment thermosensitive polymers are promising ingredients.

To geometrically structure hydrogels, various microstructuring techniques have been developed, each with specific advantages and limitations. Methods for hydrogel polymerization include chemical or physical crosslinking. While chemical crosslinking lacks control over micron scale geometry (18,24), physical crosslinking techniques, such as traditional photolithography and soft-lithography have been used to structure hydrogels on a micron scale (25,26). Additional techniques for structuring hydrogel include two-photon lithography (27,28), stereolithography (29,30), and maskless photolithography (31,32).

Here, we present an approach for generating arrays of hydrogel microstructures, that expand and contract dynamically in response to temperature modulations.We assembled a simple and cost-efficient lithographic polymerization setup for generating hydrogel structures composed of the thermoresponsive polymer NiPAAm and 4-arm-PEG-acrylate. The hydrogel microstructures exhibits robust adhesion to the substrate and undergoe reversible size changes exceeding 70% when heated above their critical temperature.

## Materials and Methods

### Hydrogel Preparation

The 4-arm-PEG hydrogel mixture was composed of 5 mM 4-arm poly (ethylene glycol)-acrylate (MW10K, Laysan Bio Inc) and 15 mM 4-Benzoybenzyl-trimethylammonium chloride (BOC Sciences) in water. The thermo-responsive gel mixture was composed of 3.75mM 4-arm poly (ethylene glycol)-acrylate, 11.25mM 4-Benzoybenzyl-trimethylammonium chloride, 1.2M Poly(N-isopropylacrylamide) (Sigma Aldrich), 1.75mM Methylenebisacrylamide (MBAm) (Thermo Scientific) and 1.75mM 2-Hydroxy-4’-(2-hydroxyethoxy)-2-methylpropiophenone (Irgacure 2959) (Sigma Aldrich) in water unless otherwise specified.

### Sample preparation

Flow chambers (Fig. 1, Fig. 2B-E, 3A,C,E, Fig. 4): Flow chambers were assembled using two glass cover slips and two spacers made of melted parafilm, resulting in an average channel height of 100 μm. A 15 μl aliquot of the liquid hydrogel mixture was introduced into the channel and polymerized via UV exposure through a photomask. For the resolution test of the setup (Fig. 2C-F), hydrogel structures were imaged immediatelly after polymerization. To characterize the temperature response of the gels, the unpolymerized gel was removed by rinsing the structures with 50μl of water prior to imaging.

**Figure 1.**
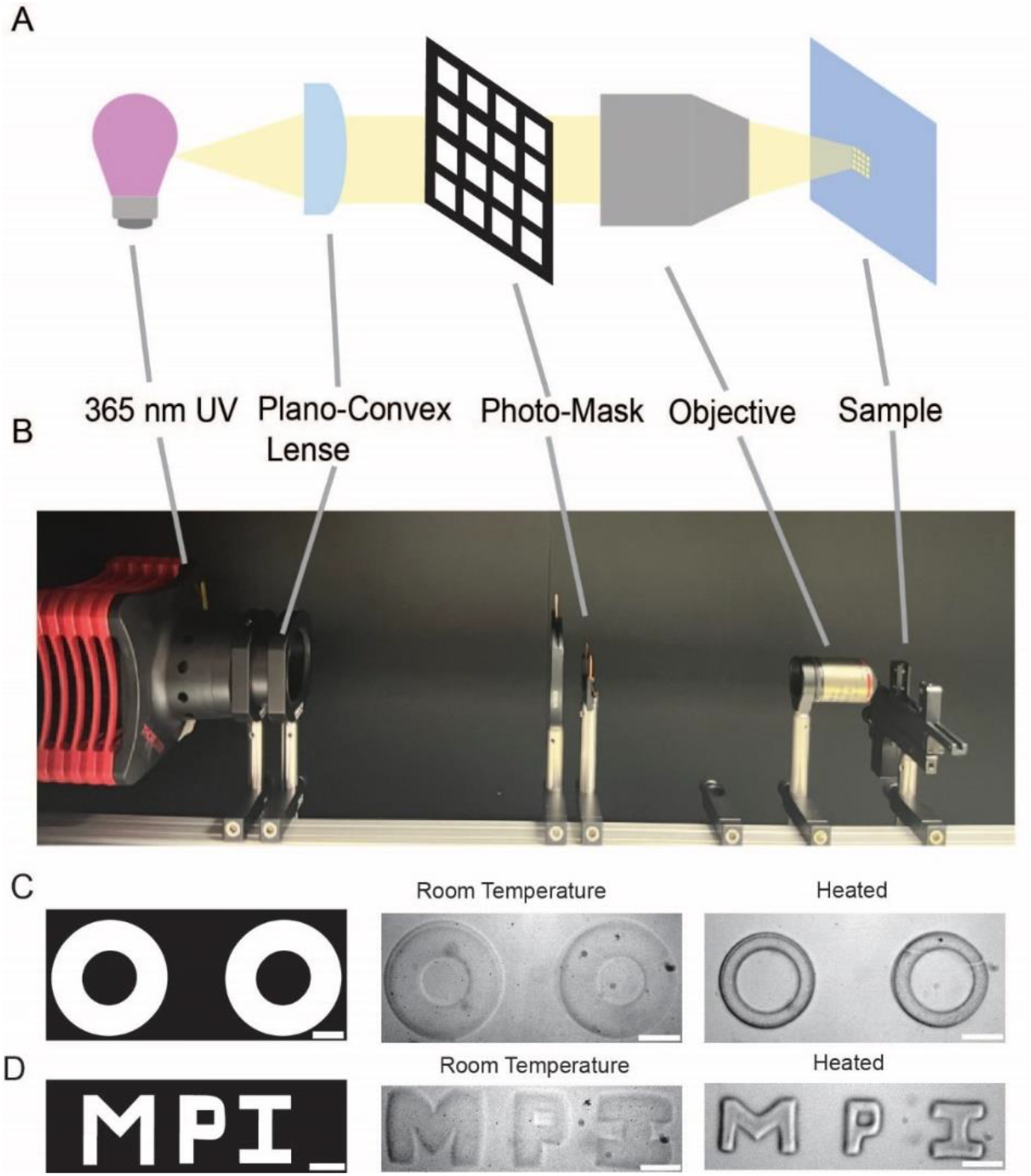
Lithographic fabrication of thermoresponsive hydrogel microstructures using a custom-made setup. A) Schematic of the lithographic projection setup. The projected light patterns have reduced dimensions compared to the photomask design. B) Photograph of the lithographic setup. (C-D) Different mask designs (left) and corresponding hydrogel microstructures. Middle: Image of polymerized hydrogel structure at room temperature. Right: Image of heated hydrogel structure. All hydrogel structures were polymerized for 10s at 391mW/cm^2^. Scale bar (design photomask): 1mm. Scale bar (microscopy images): 200μm. C) Donut-shaped structures with an outer diameter of 1.5 mm on the photomask. D) Structures forming the letters “MPI”.

**Figure 2.**
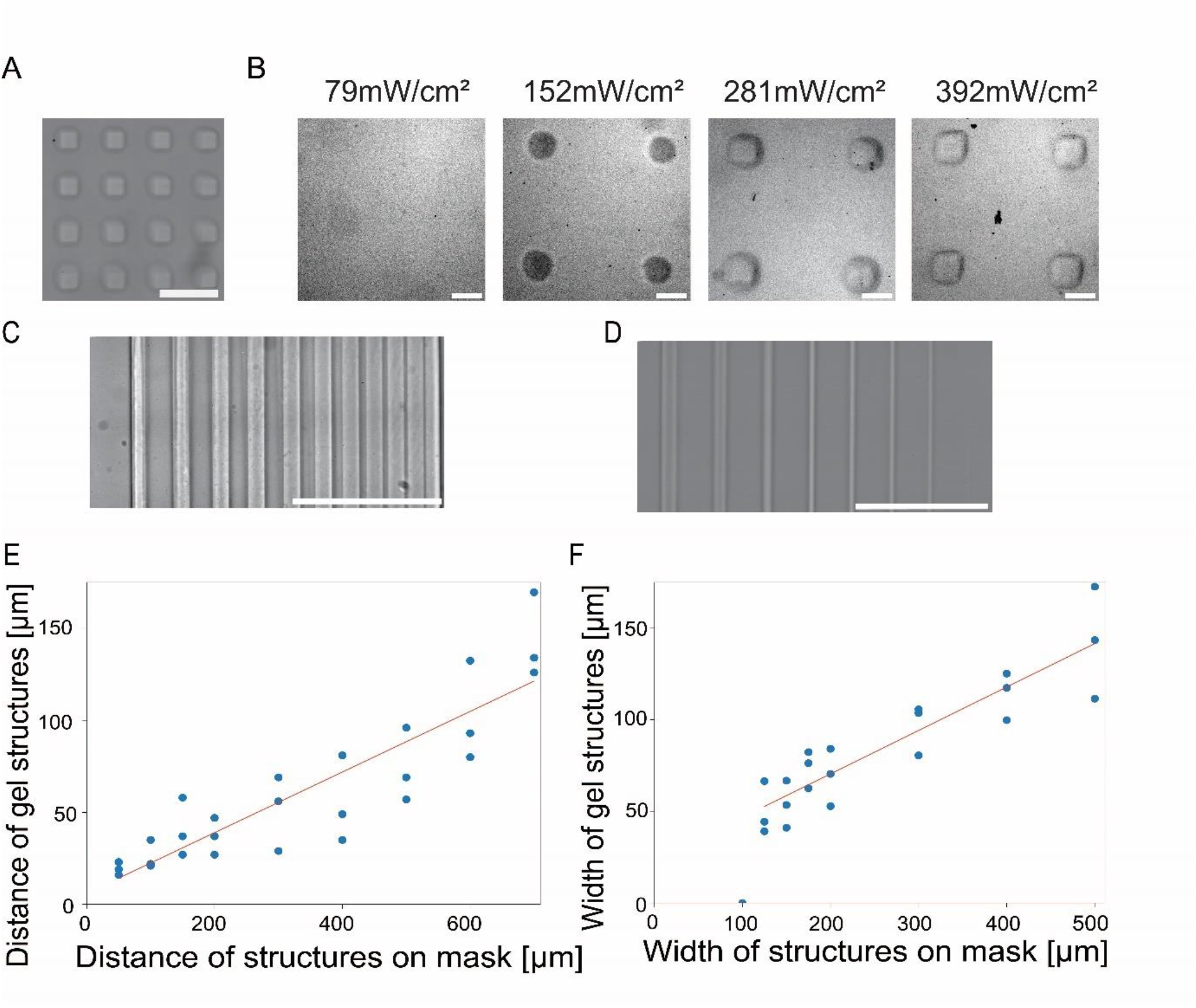
Characterization of hydrogel polymerization as a function of light intensity and template size. A) Array of hydrogel structures patterned inside a microwell using the lithographic polymerization setup. Scale bar: 500 μm B) 4-arm-PEG/NiPAAm hydrogel structures were polymerized with increasing light intensity and a constant polymerization time of 10s. Scale bar: 100 μm C) Patterned hydrogel lines with decreasing spacing between the lines. The distances between lines on the photomask ranged from 700μm to 50μm. D) Patterned hydrogel lines with decreasing line widths. The widths of the lines ranged from 500 μm to 100 μm. The 4-arm-PEG/NiPAAm hydrogel was polymerized for 10 s at 392 mW/cm^2^. E) Quantification of the distance between polymerized structures over the distance of lines on the photomask. F) Quantification of the width of the polymerized structures over the width of features on the photomask. The data were fitted with a linear function: f(x)=(0.24±0.03)*x+(23.06±6.84) μm. Independent samples: n=3

**Figure 3.**
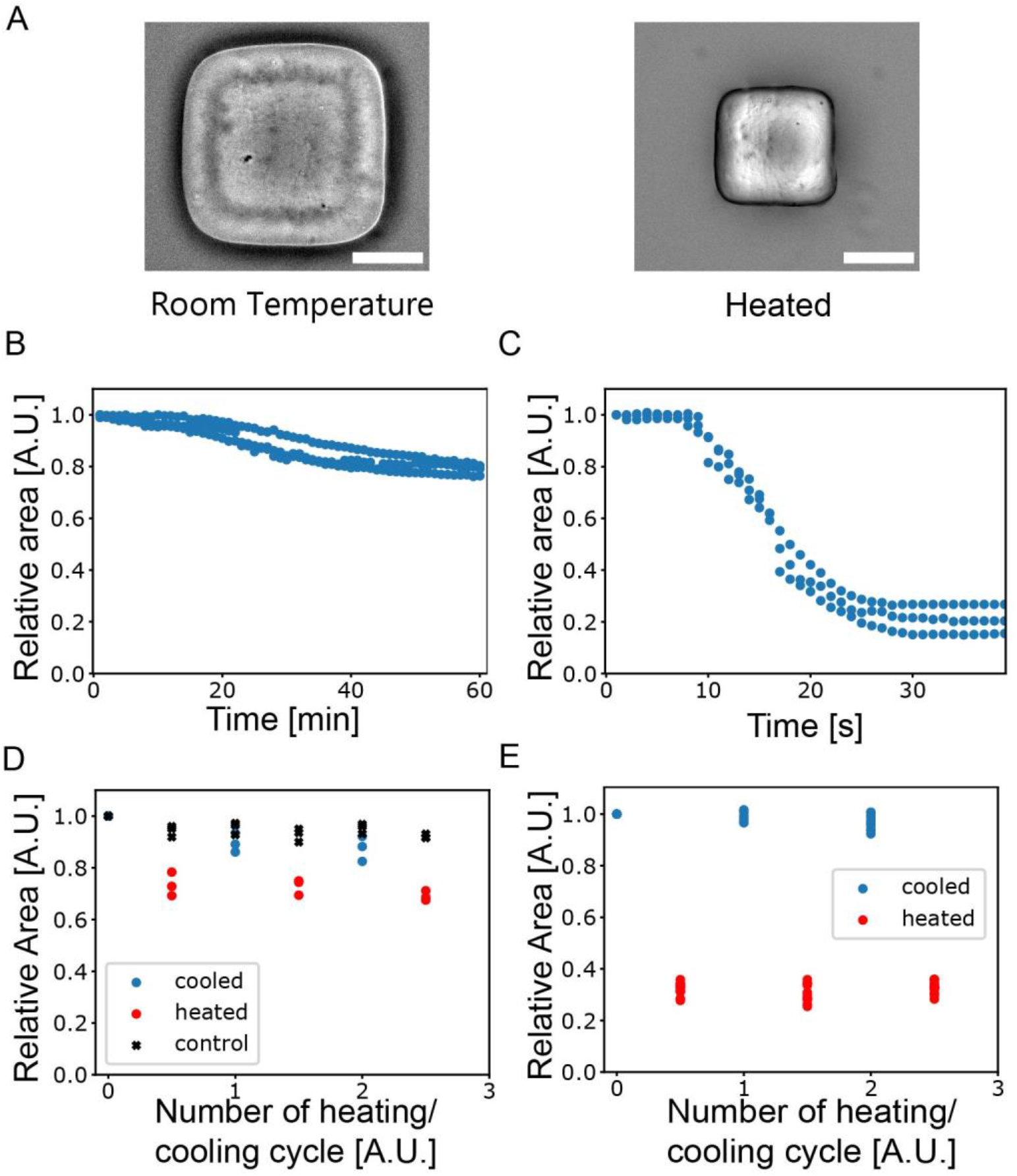
Swelling and shrinking of thermoresponsive hydrogel microstructures in response to temperature changes. A) Thermoresponsive hydrogel structures were polymerized at room temperature (left). Upon heating the hydrogel microstructures shrink (right). Scale bar: 100 μm. The samples were either heated by using a stage incubator (B, D) or by placing a glass-bottom dish filled with 70°C water on top of the sample (A, C, D). B-C) Quantification of the relative size change of the hydrogel microstructures in response to temperature variations. D-E) Repeating heating and cooling cycles were applied using a stage incubator (D) or a water bath (E) and the size of hydrogel structures was determined. A non-thermoresponsive 4-arm-PEG-acrylate hydrogel was patterned at 79 mW/mm^2^ for 7 min and used as a control (black). The 4-arm-Peg/NiPAAm hydrogel was polymerized for 10s at 392 mW/cm^2^. Independent samples: n=3

**Figure 4:**
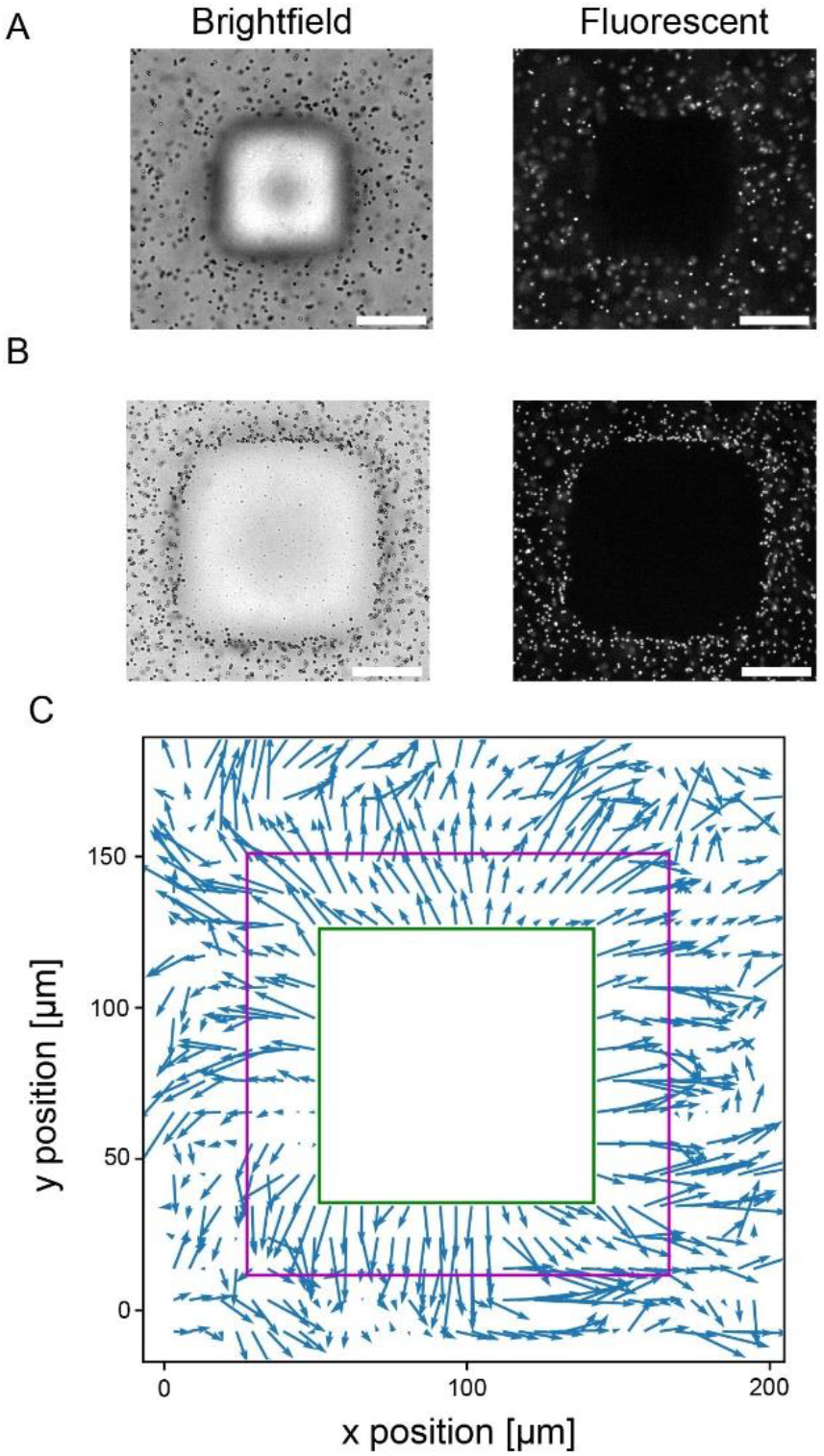
Displacement of microbeads by expanding hydrogel microstructures. A) Thermoresponsive hydrogel blocks in the contracted state, surrounded by fluorescent beads. B) The same hydrogel structures in the expanded state. Scale bar: 50 μm. C) Particle Image Velocimetry (PIV) analysis of bead movement during expansion of the hydrogel structure. The purple frame indicates the boundary of the hydrogel structure in the expanded state. The green frame indicates the size of the contracted hydrogel structure. The arrows show the directions of bead displacements away from the hydrogel structure. The 4-arm-PEG/NiPAAm hydrogel microstructure was polymerized for 10 s at 392 mW/cm^2^.

For bead displacement experiments, beads (L9529-1ml, Sigma Aldrich) were diluted 1:20 in water supplemented with 0.1% Tween-20 (Sigma Aldrich) before introducing the bead suspension into a flow chamber containing hydrogel structures.

Culture dish samples (Fig. 2A, Fig. 3B,D): A plasma-treated PDMS stencil featuring a 3 mm circular hole was placed inside a glass-bottom dish (Ibidi). Then, 5 μl of the liquid hydrogel mixture was dispensed into the resulting PDMS chamber. The chambers were closed with an PDMS sheet as lid.

Hydrogel structures were polymerized for 10 seconds at 392 mW/cm^2^, using the self-assembled projection setup, unless otherwise specified. After polymerization, the PDMS lid was removed and the hydrogel structures were rinsed with 4 ml MilliQ water.

### Light microscopy

All images were acquired using an inverted microscope (Nikon Ti2) equipped with CFI Plan Fluor 10X and CFI Plan Fluor 20X objectives. The microscopy images where analyzed using ImageJ (33).

### Temperature Modulation

Two different approaches were used to induce temperature changes. Samples in glass-bottom dishes were heated using a stage incubator (Okolab). Temperature adjustments from room temperature to 37°C typically occurred over 30-60 minutes. We refered to this heating approach as “slow heating”. Hydrogel samples within flow chambers where heated by placing a 3cm-glass-bottom dish filled with 70°C warm water on top of the samples. Cooling was achieved by removing the dish. The swelling and shrinking of thermosensitive gels occurred within less than a minute using this approach. This approach is referred to as “fast heating”.

### Optical setup for the projection of light-patterns

The optical components were purchased from Thorlabs and assembled on an optical rail system. The optical setup included a UV lamp (Thorlabs, SOLIS-365C), lenses to collimate the light beam (Thorlabs, LB1723-A and ACL25416U-A) and an iris (Thorlabs, ID50/M) to reduce the beam diameter. A 22×40mm glass coverslip with a photomask was mounted in the light path using a filter holder (Thorlabs, FH2). The sample was positioned behind an objective (Nikon, CFI Plan Fluor 4X objective) using an XY-mount (Thorlabs, XYF1/M). UV light passed through a film photo-mask (JD Photo Data) and was focussed on the sample using a 4x-objective. To ensure safety and protect the optical setup from dust, the entire system was enclosed.

### Photomask designs

Photomask patterns were designed using the software KLayout. The following patterns were used in this study:

Line structures: Stripes with widths between 100μm and 500μm were designed. Additionally, 400 μm-wide stripes with spacing between 50 μm and 700 μm were designed.

Donut structures: The donut structures had an outer diameter of 2 mm and an inner diameter of 1 mm.

Letter structures: The ‘MPI’ letters have a line width of 400 μm and a height of 1.2 mm.

### Software

The microscopy images were analyzed using Fiji (33). The Particle Image velocimetry was performed using the PIVlab (34) toolbox in Matlab (MATLAB ver. R2023a). Plots were generated using Python.

### AI

Chat GPT 3.5 (openai.com) was used to improve grammar and language.

## Results

Our goal was to engineer on a glass surface arrays of dynamic hydrogel microstructures that respond to temperature modulations. Direct photopatterning of liquid hydrogel using a transparency mask is a simple method for fabricating micrometer-scale hydrogel structures (35–37). Thus, we constructed a lithographic setup using an optical rail system (**Error! Reference source not found**.A and B). The light beam emitted by an high-power UV lamp is collimated by two lenses. Then, the beam is passed through an iris to reduce the beam diameter before reaching a transparancy photomask. The light beam passes the transparent regions of a film photomask and is focused by a 4x objective onto the liquid hydrogel material. The objective reduces the dimensions of the light patterns, resulting in hydrogel structures smaller than the photomask features.

Using the projection lithographic setup we successfully fabricated hydrogel microstructures with diverse geometries (Figure 1C-D). Initially, our attempts involved a NiPAAm-based material system composed of NiPAAm, the crosslinker MBAm, and the potoinitiator Irgacure 2959. However, during the rinsing of our hydrogel structures, the patterned hydrogel microstructures detached from the glass surface. While this effect is desireable for stop-flow lithography, where hydrogel microstructures are collected for futher applications (36,37), an assay with substrate adhesion would for instance enable micromechanical perturbation assays of adherent cells on glass bottom dishes. Thus, our next step was to improve the attachment of hydrogel microstructures to a glass surface.

Previously, various hydrogel material systems have been shown to display different adhesion properties (15,38). Therefore we hypothesized that blending 4-arm-PEG acrylate, a hydrogel material that exhibits adhesion to glass substrates, with the NiPAAm-based material system would result in a material system that maintains the thermoresponsive property of NiPAAm-based hydrogel to shink and expand while displaying increased adhesion properties.

To test this hypothesis, we prepared hydrogel mixtures of the 4-arm-PEG acrylate-based and the NiPAAm-based material system. Thereby, we varied the concentrations of the two polymer systems by using 25/75, 50/50 and 75/25 V/V ratios of 4-arm-PEG hydrogel and NiPAAm hydrogel mixture. Hydrogel structures generated with mixtures at ratios of 25/75 and 50/50 detached from the glass during the washing step, while the 75/25 4-arm-PEG/NiPAAm hydrogel remained attahed to the glass substrate. This optimized75/25 mixture consisted of 3.75mM 4-arm-PEG and 1.2M NiPAAm and was used in the following experiments (in the following referred to as 4-arm-PEG/NiPAAm).

Next, we tested the thermoresponsive behavior of the 4-arm-PEG/NiPAAm hydrogel microstructures. Using our custom-build lithographic setup, we polymerized 4-arm-PEG/NiPAAm hydrogel microstructures within a flow chamber using photomasks, featuring user defined geometries, such as donuts, and the letters “MPI” (Figure 1C-D). To induce shrinking, we placed a glass-bottom dish filled with 70°C water on top the flow chamber. The hydrogel microstructures clearly shrunk upon heating (Figure 1C-D). thereby they maintained their distinct shape and well-defined edges.

Our approach allowed to structure also larger millimeter-scale areas with arrays of hydrogel microstructures. For example, we polymerized hydrogel microstructures using a mask featuring 500×500 μm squares with a 1 mm spacing (Figure 2A). These results demonstrate the capability of our simple lithographic setup to rapidly pattern thermoresponsive hydrogels into diverse geometries, covering an area of several square millimeters.

After demonstrating that the custom-build lithographic setup can successfully polymerize 4-arm-PEG/NiPAAm hydrogel into various geometries, we investigated how increasing light intensity affects the hydrogel microstructures. To do this, we used a photomask featuring 500×500 μm squares with a 1 mm spacing and polymerized the hydrogel microstructures inside microwells for 10s at varying light intensities. We tested light intensities ranging from 79mW/cm^2^ to 392mW/cm^2^ (Figure 2B). The light intensity was measured at a distance of 10cm from the light source. The resulting polymerized structures increased with increasing light intensity and intensities below 79mW/cm^2^ failed to polymerize the hydrogel. Based on these results we selected for the subsequent experiments a polymerization condition of 392mW/mm^2^ for 10s, which polymerized well-defined hydrogel microstructures.

Next, we determined the minimal resolvable features. Initially, we used a photomask with gap sizes ranging from 50 μm to 700 μm between features (Figure 2C, E). We found that gaps of 50 μm on the mask were still resolved and resulted in a hydrogel spacing of 19±3μm. The relationship between the designed gap size on the mask and the measured structure distance followed a linear trend. To determine the smallest polymerizable structure size, we then employed a photomask featuring line widths ranging from 100 μm to 500 μm. Photomask features down to 125μm widths resulted in polymerized hydrogel structures. These hydrogel microstructures had an average width of 50±15μm. Lines with a width of 100μm on the mask did not result in hydrogel micropatterns (Figure 2D). The relationship between the line width on the mask and the resulting width of the hydrogel structures also followed a linear trend, which was described by f(x)=(0.24±0.03)*x+(23.06±6.84)μm, where x represents the line width on the mask (Figure 2E). The slope of the linear relationship represents the magnification reduction of the 4x objective.

To characterize the thermoresponsive behavior of our hydrogel microstructures, we heated hydrogel microstructures that were polymerized within a glass bottom dish using a stage incubator. The temperature was gradually increased from 22°C to 35°C over the time span of 60min. Thereby, the area of the hydrogel microstructures decreased by 21±2% (Fig. 3B). Upon cooling, the hydrogel microstructures re-expanded. To confirm that these volume changes are reproducible, we repeated the heating and cooling cycles three times per sample (Fig. 3D). The area of the hydrogel microstructures in the expanded state varied by 11.4% from the initial structure size. In the contracted state the structure area varied by 3.4% from the initial contracted state. The control 4-arm-PEG hydrogel microstructure areas varied 6.0% as compared to the original structure size and no temperature dependency was observed. Although the 4-arm-PEG/NiPAAm hydrogel microstructures underwend a clear size change compared to the control gel, the size change was relatively small compared to thermoresponsive gel objects which are not attached to surfaces (39). One reason for limiting the extend of area changes could be the friction between the expanding microstructures and the glass slide surface alongside with the slow temperature increase.

Therefore we tested whether faster temperature changes lead to larger area modulations of the hydrogel structures. To do this, we polymerized hydrogel microstructures inside a simple flow chamber. To increase the temperature more rapidly, we placed a glass bottom dish filled with 70°C water on top of the flow chamber. The areas of hydrogel microstructures shrank by 79±4% within 30 s.(Figure 3A and C). When the sample was cooled to room temperature, the structures expanded again. Repeated heating and cooling cycles resulted in size variations of 0.8% in the expanded and 2.2% in the contracted state.

The two experiments differ in some aspects: In the incubation chamber the maximum temperature was 35°C and the gel was attached to the bottom of a dish. Hydrogel microstructures in the flow chamber were attached to the bottom and top of the chamber and exposed to temperatures of up to 70°C. Previous studies have shown that even beyond the critical temperature the NiPAAm gel continues to shrink as temperature increases (40). However, despite these differences, friction that counteracts the expansion of hydrogel microstructures and slow vs. fast temperature modulations may explain our result that heating in the stage incubator led to smaller shrinkage of the hydrogel microstructures (21%), whereas the hydrogel microstructures undergo a larger area change (79%) when heated rapidly.

After characterizing the size-changing behavior of the hydrogel, we evaluated whether these size-changes influence the surrounding environment. To investigate this, we introduced fluorescent latex beads (2 μm in diameter) into a Parafilm channel containing the thermoresponsive hydrogel structures. During cooling, the gel structures expanded and pushed the beads outward (Figure 4A and B). To further analyze the displacement of the beads, we performed Particle Image Velocimetry (PIV) analysis (Fig. 4C). The results verify that the hydrogel expansions moved the beads in a directed ay outward, demonstrating that volume changes of our hydrogel can generate mechanical forces in the surrounding environment.

In summary, we engineered microstructures composed of thermoresponsive 4-arm-PEG/NiPAAm hydrogels using a simple lithographic setup. Our lithographic setup is cost-effective and leads to hydrogel structures of defined 2D geometries determined by a photomask. We demonstrated that these hydrogel microstructures exhibit a reversible size change of up to 79% in response to temperature modulations while maintaining adhesion to a glass surface. Our setup achieves a minimal structure distance of below 18μm and a minimum structure size of 50μm, offering experimental opportunities on the cellular and and tissue scale. In the future, such thermoresponsive hydrogel microstructures may be applied to induce mechanical stresses and forces and to study their effects on cellular organization.

## Acknowledgements

We thank all members of the Zieske research group for fruitful scientific discussions and for reviewing the manuscript. We also thank David Nastvogel and Marie Reischke for productive scientific discussions.

